# Time-Resolved Integrative Optical Imaging of Diffusion during Spreading Depression

**DOI:** 10.1101/648972

**Authors:** J. Hrabe, S. Hrabetova

## Abstract

An improved version of Integrative Optical Imaging method has been developed which substantially increases the time resolution of diffusion measurements. We present a theory for Time-Resolved Integrative Optical Imaging (TR-IOI) that incorporates time-dependent effective diffusion coefficient in homogeneous anisotropic media and time-dependent nonspecific linear clearance. The method was applied to measure the very fast changes in extracellular diffusion that occur during spreading depression in rat hippocampal slices. We were able to achieve time resolution of approximately one second, an improvement of at least ten times compared to the standard methods for extracellular diffusion measurement. We have found that diffusion of a small fluorescent extracellular marker (MW 3000) completely stopped during the maximum DC shift associated with the spreading depression wave, then gradually resumed over several minutes afterward. The effect of spreading depression on extracellular space is much larger than previously estimated by other methods with lower time resolution.

## INTRODUCTION

### Extracellular space

Brain cells are surrounded by extracellular space (ECS) which comprises the highly connected narrow pores filled with ionic fluid, secreted substances, and extracellular matrix. It serves multiple roles: as a reservoir for ions, a communication channel for chemical signals, and a conduit for metabolite removal. It is also a pathway for various therapeutic agents.

The typical width of ECS pores is only 40 nm (1), yet it accounts for approximately 20% of the tissue volume in a sleeping brain (2). However, the ECS is far from static. Its volume shrinks by almost half in the awake brain (3, 4) and many pathological events and brain insults, e.g., ischemia, epilepsy, brain injury, and spreading depression (SD), also lead to rapid alterations of ECS properties (5).

Molecular transport through the ECS is mediated primarily by diffusion. In addition to transporting physiologically important molecules, diffusion of experimentally introduced inert marker molecules also serves as the primary source of information about the complex ECS environment (6). In diffusion experiments, the ECS is treated as a geometrically complex porous environment characterized by two macroscopic properties: volume fraction *α* and diffusion permeability *θ* (7, 8). Volume fraction is the proportion of brain tissue volume occupied by the ECS and primarily governs concentration of molecules released into the ECS. Diffusion permeability 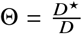, a ratio of the effective diffusion coefficient *D*^⋆^ to its value in an obstacle-free medium *D*, describes how much a diffusion-mediated process is slowed down in the ECS by obstacles represented by the cells and their various appendages (8), combined with other effects such as viscosity (9) or restricted diffusion (1). One additional parameter, *κ*, accounts for nonspecific clearance proportional to the concentration and describes removal of marker molecules over time. It should not be confused with the membrane permeability of cellular elements, assumed to be negligible for the diffusing marker macromolecules. The nonspecific clearance may occur due to, e.g., the loss of marker molecules into the blood stream *in-vivo*, or when some percentage of the fluorescent molecules stops emitting light during any fixed time interval. Penetration of marker molecules into cells would not be well described by this term unless their fluorescence was extinguished upon entry.

### Integrative Optical Imaging

Temporal resolution of currently popular ECS diffusion methods, namely the Real-Time Iontophoretic (RTI) method (10), and the Integrative Optical Imaging (IOI) method (11), is on the order of tens of seconds, which precludes their application to the conditions where fast dynamic changes occur, such as during high-frequency neuronal firing, oscillations, seizures, or SD.

In the IOI method (11, 12), a small volume of inert marker molecules labeled with a fluorescent dye is released by instantaneous pressure injection from a glass microelectrode into the brain ECS. The diffusing cloud is imaged by an epifluorescence microscope and recorded by a charge-coupled device (CCD) camera, and the image time series is evaluated to find the effective diffusion coefficient *D*^⋆^. The IOI method assumes that *D*^⋆^ is constant in time during the entire measurement, typically tens to hundreds of seconds.

The TR-IOI method improves upon IOI by allowing the *D*^⋆^ to become a function of time. Its utility was illustrated by measuring fast changes in ECS diffusion permeability during SD. The assumption is that the brain ECS environment remains homogeneous inside the sampled region. However, TR-IOI allows the medium to be diffusionally anisotropic. Surprisingly, we have found that a small dextran molecule (hydrodynamic diameter 2.8 nm) became transiently trapped in the ECS of rat hippocampal slices at the peak of SD.

### Spreading depression

Spreading depression is a transient depolarization of a group of brain cells that propagates as a slow wave (13–15). During the SD, synaptic transmission silences, electrical activity ceases (13, 14) and large changes in the concentration of ions in the ECS occur (16–19). The event lasts 1–3 minutes and brain tissue completely recovers within 10–15 minutes, after which another wave can be supported (13, 14). The SD in occipital neocortex of humans is believed to underlie visual aura associated with migraine headaches (20–22). Certain SD-like phenomena are also observed in ischemia, trauma, and during seizures (23–25).

The ECS undergoes a dramatic change during SD (26–28). The shift of ions across the cell membranes causes an osmotic imbalance resulting in gross movement of water into the cells. As the cells swell, the ECS becomes smaller and macroscopic diffusion becomes slower. The electron micrographs show that ECS in murine neocortex is often reduced to small triangular areas where three cellular elements meet. In many places, adjacent cell membranes form tight junctions (26). The decrease in the ECS volume of 40–50% in the neocortex of adult rats and cats *in vivo* and *in vitro* (17, 27, 29, 30), and 70–95% in adult rat CA1 region of hippocampus in vitro (31), was estimated from a relative change in the concentration of inert extracellular probes. The estimates of the changes in diffusion permeability, typically obtained from RTI diffusion measurements, are less numerous. A major obstacle is that the SD lasts only 1–3 minutes and the changes in the ECS are continuous and violent, hardly ever maintaining a steady state. Typical duration of one diffusion measurement using RTI, which requires steady conditions, is about two minutes. The time resolution of diffusion measurements is thus insufficient. Some investigators (28) attempted to overcome this obstacle by shortening the time of diffusion measurements to about 20 s. With this approach, they found that the ECS volume fraction changed from 0.21 to 0.09 (decrease of 57%), and the diffusion permeability for a small cation tetramethylammonium (TMA, hydrodynamic diameter 0.5 nm) decreased from 0.37 to 0.25 during the SD in adult rat neocortex in vivo. The TR-IOI technique achieved temporal resolution of approximately 1 s during the entire SD events, an improvement of about 20 times.

## THEORY

Let concentration *c*(**r**, *t*) of some extracellular marker molecule be a function of position in space **r** = (*x*_1_, *x*_2_, *x*_3_) and time *t*. In a geometrically complex ECS, all the diffusion parameters are defined as volume-averaged local quantities over a sufficiently large sampling volume (8). Depending on the molecular weight and the average size of cellular elements (typically units of micrometers), it usually takes about a second or two after releasing macromolecules from any point source inside the ECS to establish such averaging. The concentration is defined in terms of the total tissue volume rather than the ECS volume because optical methods such as the IOI cannot distinguish between different tissue compartments. We assume that a homogeneous but anisotropic environment with time-dependent diffusion characteristics can be described by an effective diffusion tensor

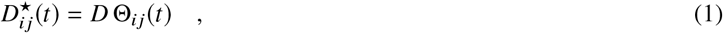

where *D* is the scalar free diffusion constant and Θ_*ij*_ is the diffusion permeability tensor. Both indices run from 1 to 3. In an environment where nonspecific loss of diffusing substance is proportional to the concentration, we also introduce linear time-dependent clearance *κ*(*t*), which is also assumed to be homogeneous. The diffusion equation in this environment is

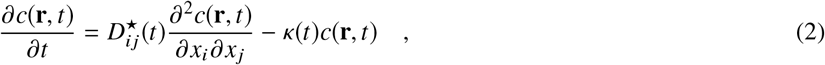

where we used Einstein’s summation convention over any index appearing exactly twice 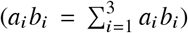. Equation 2 expresses mass conservation, providing the diffusion flow **J** obeys Fick’s law

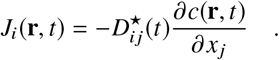

The initial concentration at time *t*_0_ is represented by a function *c*(**r**, *t*_0_).

To extract the effective diffusion as a function of time from the experimentally accessible quantities, we will first obtain solution of Eq. 2 in the form of a spatial convolution of the initial condition with the concentration resulting from a point source. It will be shown that the effective diffusion tensor at any time can be obtained from the 3D concentration cloud by calculating the time derivative of the tensor of its second moments. This can be done regardless of the initial condition or the nonspecific clearance. While this would be fairly straightforward if the 3D data were available, the traditional IOI setup acquires images in a single focal plane using an ordinary epifluorescence microscope. We therefore proceed to take such experimental situation into account and also examine the effect of the microscope’s Point-Spread Function (PSF). In this case, only the planar elements of the effective diffusion tensor can be extracted and certain requirements have to be imposed on the initial distribution of molecules. We will show that if such conditions are met, the focal plane elements of the effective diffusion tensor can still be extracted from the recorded images of the diffusion cloud as time derivatives of their second moments. Finally, it will be demonstrated that the TR-IOI theory contains the previously published IOI theory as a special case, one with a constant effective diffusion and a point source initial condition.

Equation 2 with its initial condition can be solved in the Fourier domain. Fourier transform of *c*(**r**, *t*) with respect to the three spatial coordinates is defined as

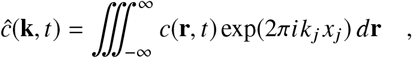

and the inverse as

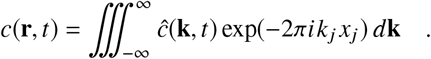

The Fourier transform turns Eq. 2 into

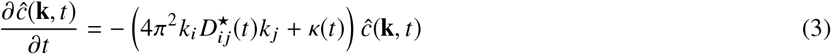

with the initial condition 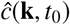.

Eq. 3 can be solved by direct integration:

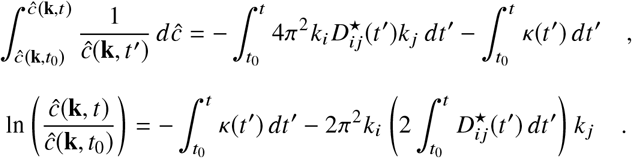

The solution can therefore be written as

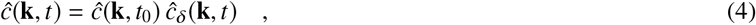

where

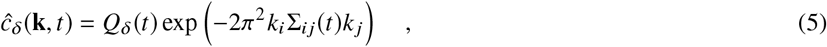

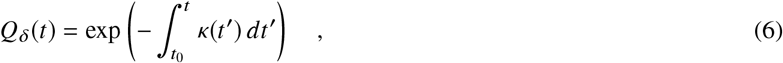

and

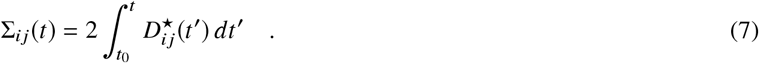

When the initial condition is Dirac’s *δ*-function *δ*(**r**), its Fourier transform is unity. The inverse Fourier transform *c*_*δ*_(**r**, *t*) of 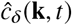 therefore describes the diffusion cloud initiated by the point source at time *t* = *t*_0_:

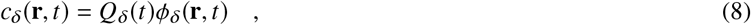

where

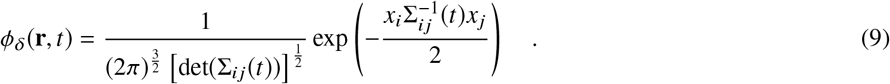

The concentration cloud *c*_*δ*_(**r**, *t*) is shaped as a 3D Gaussian distribution with variance matrix Σ_*ij*_(*t*), and the amount of diffusing substance contained in it diminishes as *Q*_*δ*_(*t*) from its initial value of *Q*_*δ*_(*t*_0_) = 1.

Multiplication in the Fourier domain corresponds to a convolution in the spatial domain. Based on Eq. 4, the concentration distribution following an arbitrary initial condition can therefore be written as

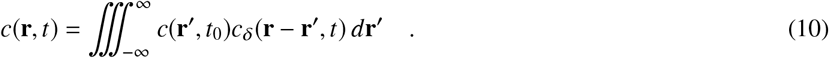

The total amount of diffusing substance is initially 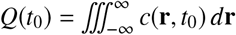 and changes with time as

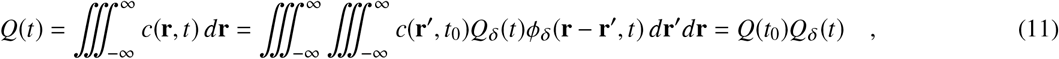

where *Q*_*δ*_(*t*) is substituted from Eq. 6. If a 3D measurement of concentration in time was available, the clearance *κ*(*t*) could be computed from Eqs. 11 and 6:

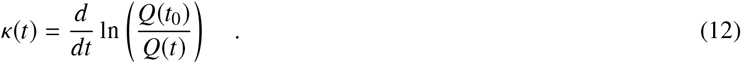

Since the substance loss is assumed homogeneous in space, the effect of nonzero clearance simply amounts to a global scaling of amplitude.

If it is possible to normalize the measured 3D concentration by the total amount *Q*(*t*) of the diffusing substance at every time, a probability density function

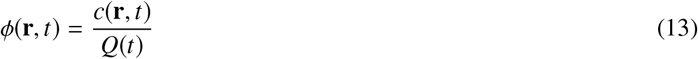

can be constructed and the tensor of its second moments *μ*_*ij*_(*t*) computed:

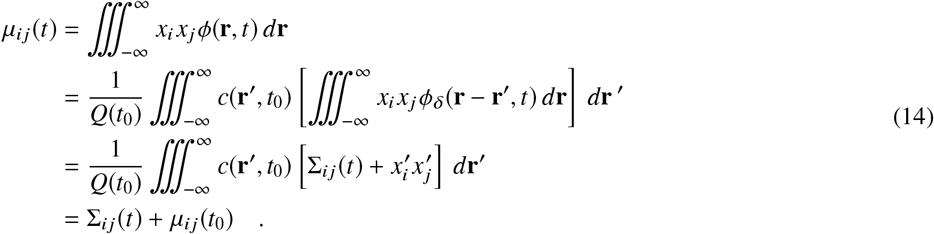

The components of the effective diffusion tensor could now easily be extracted with the help of Eq. 7 as time derivatives of these moments:

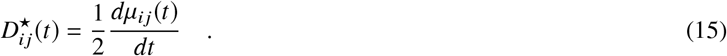

Note that Eq. 15 is valid for an arbitrarily shaped initial condition. Unfortunately, a complete 3D measurement of the concentration is not usually available. Common experimental setups, based on traditional, confocal or two-photon microscopy, record 2D images at a single focal plane. Such systems appear to be imaging a virtual object, constructed from the true object by a convolution with the PSF *S*(**r**) of the system. The effective width of the PSF limits the system resolution. Using the approximation for *S*(**r**) suggested by (11, 12), we assume that the horizontal resolution is higher than the corresponding size of the recorded image pixel and the horizontal PSF effect can thus be ignored. The PSF approximation in the object space is then

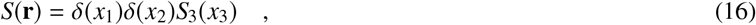

where *S*_3_(*x*_3_) causes blurring along the optical axis. The exact functional form of *S*_3_(*x*_3_) is not important as long as the approximation given by Eq. 16 remains valid.

If the PSF is very sharp (*S*(**r**) = *δ*(**r**)), as is the case in confocal or two-photon setups, the imaging system simply records a 2D image proportional to the concentrations *c*(*x*_1_, *x*_2_, *x*_3_ = *z*, *t*) in the plane of focus *x*_3_ = *z*. In traditional microscopy setup however, it instead appears to detect signal originating from concentration

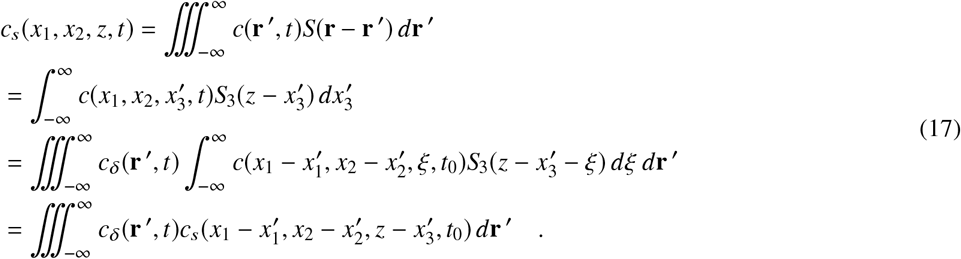

It can be seen that the effect of microscope’s PSF results in a simple modification (blurring along the *x*_3_ axis) of the initial condition *c*(**r**, *t*_0_) to *c*_*s*_ (**r**, *t*_0_). After this modification, we can consider the microscope to perfectly follow the rules of geometrical optics, magnifying the image by a constant factor *M* and amplifying the image signal by another constant amplitude factor *A*. A single in-focus plane *x*_3_ = *z* through *c*_*s*_ is imaged.

The IOI and TR-IOI methods acquire data in a single focal plane and therefore cannot accommodate an arbitrary orientation of the effective diffusion tensor axes. That would require additional measurements acquired in two other (independently oriented) planes. In general, diffusion along six independent directions would have to be evaluated to extract the six components of the symmetric diffusion tensor. In practice, it is often possible to make one of the principle axes of the effective diffusion tensor parallel to the optical axis of the imaging system to obtain the two remaining principal diffusivities, so this is usually not a serious limitation. Consequently, the math can be simplified by working in a coordinate system where the *x*_3_-axis coincides with one of the principal axes of the diffusion tensor. We then have

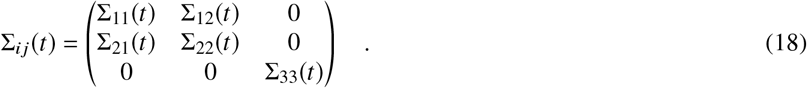

Assuming that a 2D section at *x*_3_ = *z* of the concentration cloud elicited by the blurred initial condition *c*_*s*_ (**r**, *t*_0_) represents all the information that is available to us, let us extract as much as possible from it. The image intensity *I*(*x*_1_, *x*_2_, *t*) expressed in the object coordinates after constant amplification *A* is

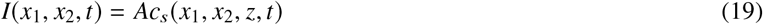

For the integrated total image intensity *Q*_*I*_ (*t*) we obtain

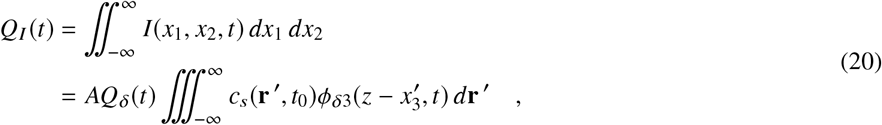

where

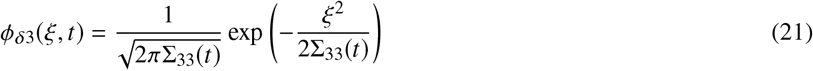

is the “vertical” portion of the Gaussian distribution *ϕ*_*δ*_. In contrast to the 3D measurement, the time dependency is no longer fully determined by the clearance term *Q*_*δ*_(*t*).

Having obtained the normalization factor *Q*_*I*_ (*t*), we can calculate the second moment *μ*_*JK*_ (*t*) of the image. Capitalized indices are introduced to distinguish their 2D range (*J*, *K* = 1, 2).

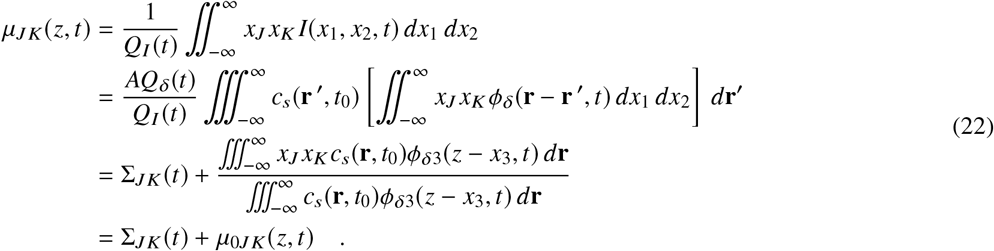

Our ability to proceed similarly as in Eq. 15 and extract the effective diffusion tensor components 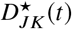 from the time derivative

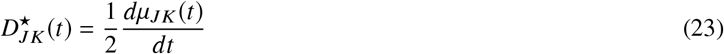

depends on the “nuisance term” *μ*_0*JK*_ (*z*, *t*), which unfortunately depends on time. This problem is not unique to the TR-IOI. The same reasoning is valid even when the diffusion tensor is constant, as it is assumed to be in the traditional IOI method. However, if the initial condition *c*(**r**, *t*_0_) is a separable function in its spatial variables, then *c*_*s*_ must be separable as well; the time-dependent terms in the numerator and the denominator of *μ*_0*JK*_ cancel out, and *μ*_0*JK*_ becomes constant in time. The most common form of the initial condition assumed also in (11), the 3D Gaussian, falls into this category. Another interesting possibility is that an objective with low numerical aperture effectively projects the entire diffusion cloud along the vertical direction. The *c*_*s*_ (**r**, *t*_0_) then does not depend significantly on *z* and the nuisance term again becomes constant.

It is easy to verify that the TR-IOI theory contains the IOI as a special case. When the diffusion cloud originates from a 3D Gaussian initial condition in an environment characterized by constant and isotropic effective diffusion tensor, Eq. 22 simplifies to

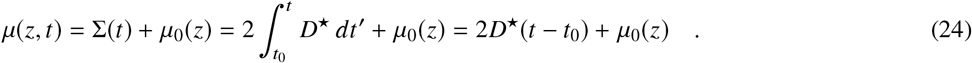

This is equivalent to the IOI equation derived in (11), except that the authors extrapolate the experiment back in time so that *t*_0_ becomes the time when *μ*_0_(*z*) would have reached 0 because the initial Gaussian would have shrunk to a *δ*-function. In IOI, the constant effective diffusion coefficient *D*^⋆^ can be extracted by linear regression.

## MATERIALS AND METHODS

### Rat brain slices

All experiments were conducted at NYU School of Medicine and SUNY Downstate Medical Center in accordance with NIH guidelines and local Institutional Animal Care and Use Committee regulations. Brain slices were prepared from adult Sprague-Dawley female rats (200–250 g) as described previously (32, 33). The animals were anesthetized by intraperitoneal injection of sodium pentobarbital (65 mg/kg) and decapitated. The brains were extracted and cooled down with ice-cold artificial cerebrospinal fluid (ACSF; 124 mM NaCl, 5 mM KCl, 26 mM NaHCO_3_, 1.25 mM NaH_2_PO_4_, 10 mM D-glucose, 1.3 mM MgCl_2_, and 1.5 mM CaCl_2_). For TMA diffusion experiments, 0.5 mM TMA chloride was added to the ACSF to provide a calibration standard. The osmolality of the ACSF was determined with a freezing point depression osmometer (Osmette A #5002; Precision Systems Inc., Natick, MA). A typical value of normal ACSF was 300 mosmol/kg.

Coronal sections 400 *μ*m thick, containing hippocampus, were cut on a vibrating blade microtome (Leica VT 1000 S; Leica Microsystems, Wetzlar, Germany). The slices were incubated for at least 1 hour, submerged at room temperature in the ACSF gassed with a mixture of 95% O_2_ and 5% CO_2_ to stabilize the pH at 7.4.

### Induction and recording of SD

For SD diffusion measurements, one slice at a time was transferred to a submersion tissue chamber (Warner model RC-27L; Harvard Apparatus, Holliston, MA) and superfused with ACSF at a flow rate of 2.0 mL/min. To facilitate elicitation of the SD, 50% of NaCl in the ACSF was replaced with Na propionate (30, 34, 35). The temperature of the flowing ACSF was maintained at 34° C using an in-line heater (Warner model SH-27A; Harvard Apparatus) in combination with a chamber heating system connected to a dual automatic temperature controller (Warner model TC-344B; Harvard Apparatus).

All recordings were made from the stratum radiatum in the CA1 region of the hippocampus (Fig. 1). Two glass double-barrel micropipettes were placed in the tissue at the depth of 200 *μ*m from the upper slice surface. The first double-barrel micropipette was placed on the Shaffer collaterals in the area between CA3 and CA1 regions, the induction site of the SD. For the SD induction, KCl (1 M) was released from the first barrel of this micropipette by a brief pressure pulse of nitrogen. The second barrel was filled with NaCl (150 mM) and was used to confirm that a negative DC shift reflecting local depolarization occurred at the injection site, giving a clear indication that sufficient KCl was released from the micropipette to depolarize a population of cells. The second double-barrel micropipette was placed in the stratum radiatum of the CA1 region, the recording site of the diffusion and the SD. For the diffusion measurement, dextran tagged with Texas Red was released from one barrel of a micropipette by a brief pressure pulse of nitrogen; a series of images were taken with the CCD camera. The second barrel detected the DC potential to confirm that SD did indeed arrive at the recording site, and to establish at what time this occurred.

**Figure 1:**
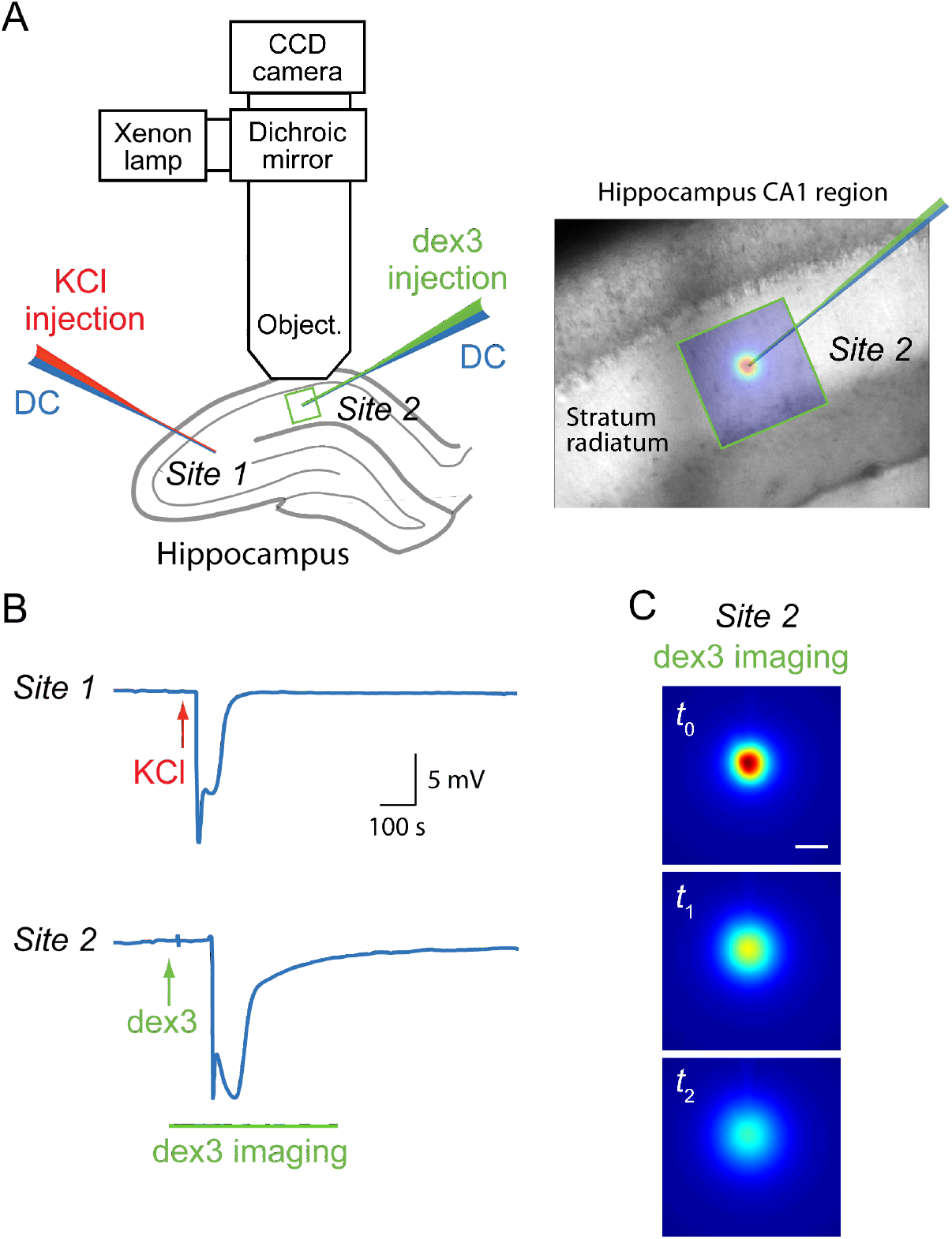
Simultaneous recording of the SD and the extracellular diffusion of dextran (MW 3000; dex3). **A.** Left: Placement of two double-barrel micropipettes in hippocampus and position of an epifluorescence microscope objective (Object). SD was induced (Site 1) by KCl injection and recorded (DC) both locally (Site 1) and at the site of dex3 diffusion (Site 2). Right: A micropipette and a fluorescent image of dex3 diffusion cloud in the stratum radiatum (Site 2) superimposed on a bright field image of the CA1 hippocampal region. **B.** Experimental protocol: 1) recordings of stable DC potentials at Site 1 and Site 2; 2) imaging of dex3 diffusion starts at Site 2; 3) SD is induced at Site 1; 4) SD arrives at Site 2 and impacts diffusion of dex3, which is recorded as a time series of fluorescent images (**C**). The scale bar is 100 μm.

The microelectrodes were held in two separate robotic micromanipulators (MP 285; Sutter Instrument Co., Novato, CA). Brief pressure pulses of compressed nitrogen were delivered by electronically controlled pressure injection unit (Picospritzer; Parken Hannifin Corp., Fairfield, NJ). The DC potentials in both microelectrodes were amplified by a CyberAmp 320 amplifier (Axon Instruments, Inc., Union City, CA), low-pass filtered (6 Hz), and plotted by a dual-channel chart recorder.

### Image acquisition

The TR-IOI and the IOI experimental setups are the same. Fluorescent diffusion marker is pressure injected from a glass micropipette and its spatial distribution is recorded by a CCD camera. The micropipette is manufactured with a fine tip of about 2 μm diameter. Short injection times (50–150 ms) aim to inject very small volumes, estimates to be less than 1 nL (11). We used dextran marker molecule (dex3, MW 3000) labeled with Texas Red (catalog number D-3329; ThermoFisher Scientific, Waltham, MA) dissolved in 150 mM NaCl to give a final dextran concentration of 1 mM. Fluorescent dextran was then pressure-injected into the diluted agarose gel (for testing or reference purposes) or the brain slice.

The excitation light from a 75 W xenon bulb mounted on the epifluorescence port of the Olympus BX50WI compound microscope (Olympus America, Melville, NY) was directed to the specimen using a dichroic mirror through a Texas Red filter set. The microscope was equipped with a water-immersion objective (Olympus, UMPlanFl, 10 ×, NA 0.3). The light emitted by the excited diffusing molecules was detected and imaged at time intervals of 1–2 s by a CCD camera (model CH350, Photometrics, Tuscon, AZ) attached to the microscope. Typically, 200–300 images were recorded over 5–10 min. Both the injections and the image acquisitions were controlled programmatically from an acquisition computer.

### TR-IOI data analysis

The diffusion analysis of the acquired sets of images was performed on a Linux workstation using an in-house program written in IDL language (Harris Geospatial Solutions, Broomfield, CO). First, 1–3 background images recorded before the dye injection were averaged and subtracted from all the diffusion images. This helped to reduce the influence of light scattering from the dye in the injection micropipette. In addition, a binary mask was constructed to exclude the micropipette and its immediate vicinity from the analysis.

Similarly to the traditional IOI data analysis, it was assumed that the diffusion cloud originated from a Gaussian initial condition. Because no anisotropy was detected during preliminary analysis, the tissue environment was assumed to be isotropic. The image of the diffusion cloud at every time point was therefore fitted by an isotropic 2D Gaussian function to obtain the second moment *μ* = *μ*(*t*) (equivalent to the variance of a symmetric 2D Gaussian distribution) as a function of time. The fitting procedure employed Marquardt least-squares estimator (36) and took advantage of the known analytical derivatives with respect to the fitting parameters.

Based on Eq. 23, the scalar effective diffusion coefficient *D*^⋆^ was estimated from the time series of the second moments *μ*(*t*). Because numerical derivatives are equivalent to high-pass filters and increase noise, the time series was first smoothed by approximating it with a linear combination of Legendre polynomials of an order up to 12. Because these polynomials are orthogonal on interval < −1, 1 >, the time series was mapped onto this interval, fitted using a singular value decomposition least-squares routine (37) and rescaled to the original time span. The fitting procedure was visualized to check for undesirable oscillations or other problems. The time series of effective diffusion coefficients *D*^⋆^(*t*) was then extracted by taking a derivative of the fitted polynomial. This method proved superior to a simpler top-hat smoothing of the *μ*(*t*) time series, which was also tested but is not shown. When diffusion coefficient was expected to be constant (e.g., in the agarose or in brain before the SD was induced), we simply reduced the Legendre polynomial to the order of 2 and extracted the slope of this linear fit.

### Real-Time Iontophoretic method

In the RTI method (10), a small extracellular probe, in our case the cation TMA (MW 74), was released from an iontophoretic microelectrode and detected by an ion-selective microelectrode positioned 120 μm away. We followed the protocol described in detail in (38) to perform these measurements in diluted agarose gel or in the brain slices, and to analyze the diffusion curves. Diffusion measurements in diluted agarose gel provided the transport number *n*_*t*_ of the iontophoretic microelectrode, i.e., the fraction of iontophoretic current carrying the TMA cations, and the free diffusion coefficient *D*. The values of *n*_*t*_ and *D* were utilized when analyzing the measurements in brain to yield the ECS volume fraction *α*, the effective diffusion coefficient *D*^⋆^, and the nonspecific clearance *κ*.

## RESULTS

### Validation of TR-IOI analysis

We began by validating the TR-IOI analysis in static conditions. The TR-IOI analysis quantified diffusion of dex3 in agarose gel and in brain slices under normal steady conditions. The results obtained with TR-IOI were in excellent agreement with those obtained by the traditional IOI analysis (Table 1).

**Table 1:** Validation of TR-IOI analysis

| Medium | Parameter<sup>a</sup> | IOI Analysis | TR-IOI Analysis | $n^b$ |
| --- | --- | --- | --- | --- |
| Agarose | $D$ | $22.0 \pm 1.1$ | $21.8 \pm 1.3$ | 5 |
| Brain | $D^*$ | $9.6 \pm 0.7$ | $9.8 \pm 0.9$ | 6 |

<sup>a</sup> [ $10^{-7} \text{ cm}^2/\text{s}$ ]; obtained at 34° C; mean ± sd.
<sup>b</sup> Number of measurements.

### Effect of propionate

Propionate was added to ACSF to facilitate SD induction, as described in Methods. However, we noticed that the diffusion permeability for dex3 was lower in the ACSF containing propionate (*θ* = 0.26) than without it (*θ* = 0.42 reported in (33)). We therefore employed RTI method to quantify the effect of partial chloride substitution on ECS volume fraction *α* and diffusion permeability *θ* for TMA. The results from 5 slices obtained from 5 animals are summarized in Fig. 2. When ACSF contained the propionate, both the ECS volume fraction and the diffusion permeability significantly decreased. Following the wash-out of propionate ACSF with regular ACSF, the diffusion permeability returned to its control value and the ECS volume fraction even exceeded it. Statistically significant differences were determined with ANOVA (*p* < 0.001 for ANOVA, *p* < 0.05 for post hoc pairwise comparison tests) and are marked with an asterisk in Fig. 2. There were also statistically significant differences in the non-specific clearance *κ* among the three groups (control: 0.013 ± 0.001, propionate: 0.014 ± 0.01, wash-out: 0.010 ± 0.001).

**Figure 2:**
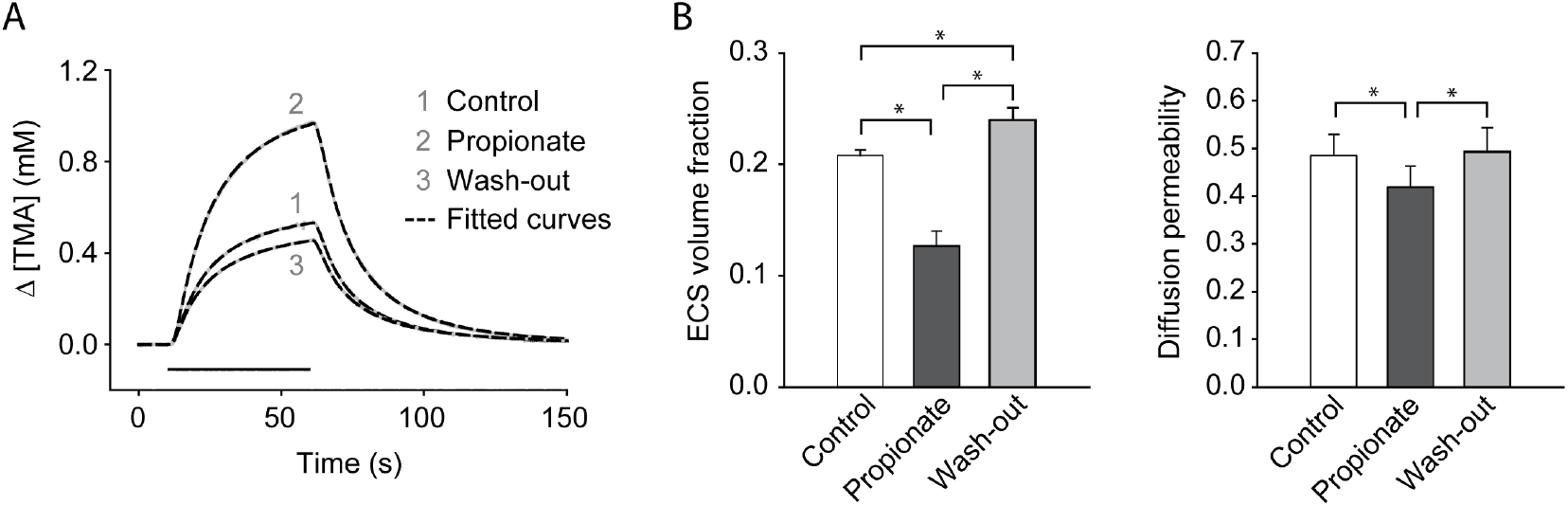
RTI study of the effect of propionate on ECS volume fraction *α* and diffusion permeability *θ*. **A.** Representative examples of TMA diffusion records in CA1 stratum radiatum under control conditions, after application of ACSF containing Na-propionate, and during wash-out. Recorded TMA curves (gray lines) with the fitted theoretical curves (dashed black lines) are shown. The ECS volume fraction *α* was 0.21 for the control record, 0.12 for the propionate record and 0.25 for the wash-out record; diffusion permeabilities *θ* were 0.490, 0.407, and 0.485; non-specific clearances *κ* were 0.012 s^−1^, 0.016 s^−1^, and 0.009 s^−1^. For all three records, the microelectrodes were spaced at 120 *μ*m and the transport number *n*_*t*_ was 0.29. The horizontal bar marks the TMA release. **B.** Summary of experimental results in CA1 stratum radiatum under three experimental conditions. Data are displayed as mean ± sd. Statistically significant differences are marked with asterisks. See text for details.

### Diffusion of dextran during SD

A representative experiment is shown in Fig. 3. Similar results were obtained in 9 SDs induced in 6 slices prepared from 3 animals. The diffusion of dex3 was recorded for about 1 minute under control conditions (*θ* = 0.26 ± 0.05). When the SD reached the diffusion cloud, a dramatic three-phase change occurred. In the first phase, lasting about 10 s, the diffusion coefficient decreased. In the second phase, lasting about 60 s, the molecules stopped moving entirely. In the third phase, the molecules started to move again but with the effective diffusion coefficient much lower (t-test, *p* < 0.001) than before the SD event (*θ* = 0.11 ± 0.08). In one experiment, another diffusion measurement was taken one hour after the SD event, suggesting a complete recovery (*θ* = 0.24).

**Figure 3:**
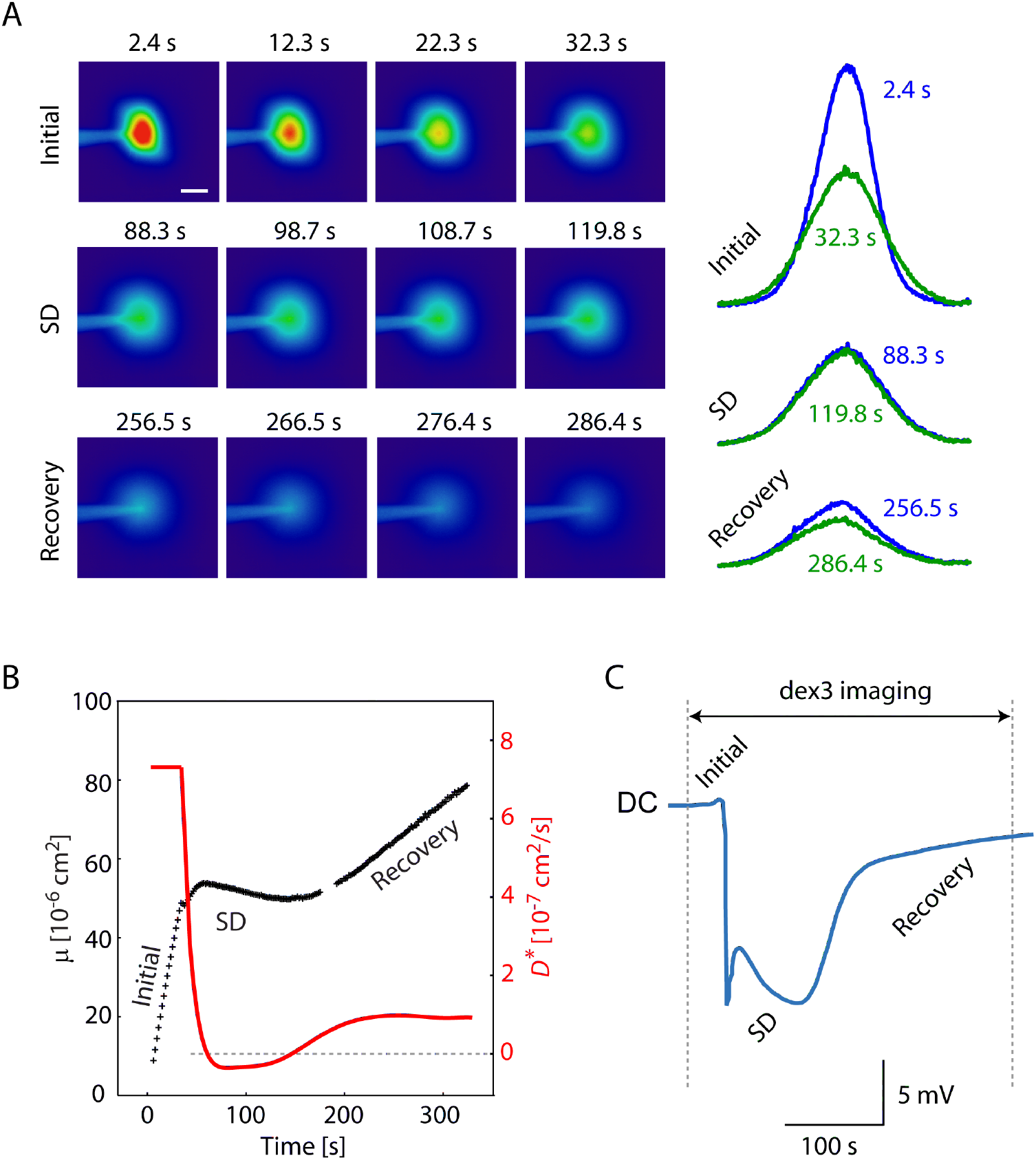
Extracellular diffusion of dex3 is transiently halted during SD. **A.** Three sequences of dex3 diffusion images taken prior to SD (Initial), at the peak of SD (SD), and after SD (Recovery). Intensity profiles extracted at two time points from these images are shown on the right. The scale bar is 100 μm. **B.** Progression of the diffusion cloud is characterized by *μ* = *μ*(*t*) and *D*^⋆^ = *D*^⋆^(*t*). During SD, *μ*(*t*) is no longer linear in time; its derivative determines the *D*^⋆^(*t*) at any given time. At the peak of SD, the spread of dex3 is transiently halted. The dotted line marks *D*^⋆^ = 0. **C.** The corresponding DC record.

Interestingly, the effective diffusion appeared to be transiently negative in the example shown. This was not a consistent phenomenon, so the cause is unclear. One possibility is that some molecules were already trapped while others still found their way through the tissue. This could temporarily disturb the Gaussian shape of the cloud and the fit for variance. It may also just be a random variation in this particular experiment.

## DISCUSSION

In Eqs. 22 and 23, we have obtained a result useful for employing the TR-IOI modification of the IOI method. The experimentally accessible quantity is *μ*_*JK*_(*t*), while its time derivative yields the corresponding components of the effective diffusion tensor 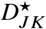 and diffusion permeability tensor *θ*_*JK*_. However, for both the IOI and the TR-IOI, it is required that the additive term *μ*_0*JK*_(*t*) related to the initial condition is constant in time. As was already mentioned, this is guaranteed for an initial condition separable in Cartesian spatial variables or for a 2D projection of a 3D diffusion cloud. Another possibility occurs if the blurred initial condition can be considered negligibly small beyond a sphere of some finite radius *ρ*. For long enough times then, namely when Σ_33_(*t*) ≫ *ρ*^2^, the *ϕ*_*δ*3_ becomes almost constant inside this sphere, leading again to *μ*_0*JK*_ constant in time. This is not surprising because after a sufficiently long interval of time, the detailed shape of the initial condition with limited spatial extent becomes irrelevant; the diffusion process “forgets its past”. However, non-separable initial condition may masquerade as initially time-dependent diffusion tensor and could be misinterpreted as such, or as an indicator of anomalous diffusion. It is therefore important to keep the injection volumes small. Numerical model using a sharp PSF indicates that, starting from initially homogeneous sphere with a diameter of 100 μm, diffusion of dex3 would resemble normal diffusion quite early; the error in *D*^⋆^ would become less than 1% after 10 s from the injection. This problem is therefore not likely to be a serious one under normal experimental conditions.

Another requirement for the TR-IOI analysis is that the principal axes of the effective diffusion tensor do not rotate out of the focal plane during the measurement. The focal plane sampling used in both the IOI or the TR-IOI cannot obtain effective diffusion tensor in arbitrary orientation; that would require information along at least six independent directions, similarly to the diffusion tensor imaging method using magnetic resonance (39). In practice, the principal axes can usually be inferred from the underlying geometry, e.g., of parallel fibers (40). During SD, the wave of disturbance propagates with a velocity of about 3 mm/min across the field of view of about 0.5 mm, taking about 10 s to cross it. This could induce a transient anisotropy in the diffusion but we did not detect any such phenomenon.

One potential limitation of TR-IOI is the decreased signal-to-noise ratio (SNR) of the effective diffusion tensor, because it is necessarily extracted from a shorter image sequence. This is the price paid for estimating *D*^⋆^ as a function of time rather than a constant. One partial remedy is to decrease the image acquisition interval as much as possible. Although it may also cause the photobleaching effect, the TR-IOI is fairly robust in this respect, similarly to IOI, because it depends only on calculation of the second moment of the image signal, irrespective of a gradual loss of signal over time. Another helpful remedy is using the whole image instead of the extraction of several profiles described in (11). We have successfully done so in this study. Both these approaches increase the number of measured or processed signal samples, which results in higher SNR. Naturally, there will always be imposed a trade-off between the TR-IOI time resolution and the SNR. Using longer interval of an acquired time series to extract the diffusion tensor at any given time will result in higher SNR but at the same time reduce the time resolution.

Previous studies reported that diffusion permeability for a small extracellular marker TMA (hydrodynamic diameter 0.5 nm) decreased by 32% during SD (28). However, these experiments utilized RTI method that requires steady state conditions while taking a measurement (i.e., at least 20 s or more) and therefore does not sufficiently resolve fast dynamic changes during SD. The TR-IOI techniques provides higher temporal resolution than the RTI method. For SD experiments reported here, we used temporal resolution 1–2 s and observed dramatic changes in diffusion permeability for dex3, including temporary entrapment at the peak of SD. The TR-IOI technique can be utilized to study extracellular diffusion during other fast dynamic events, such as oscillations and epilepsy, and its temporal resolution could be further enhanced. TR-IOI also allows to study diffusion of a large variety of small and large molecules, as long as they are fluorescent or fluorescently-labeled.

It has been recognized that replacing a part of extracellular chloride with larger anions facilitates SD induction (41) but the mechanisms remained unknown. Several mechanisms were considered, including GABA-induced depolarization, altered anion transport, pH-related effects, and changes in coupling by gap junctions (34, 35). We have found that replacing 50% on NaCl with Na propionate lead to a significant reduction of ECS volume, which facilitated SD induction. ECS volume reduction is known to similarly facilitate the epileptiform activity (42).

One interesting possibility is to combine our results with the restricted diffusion theory (43, 44), which was previously used to estimate the brain intercellular spacing (1). For a macromolecule with hydrodynamic diameter *d* diffusing within sheet-like pores of width w, the restricted diffusion theory predicts diffusion permeability

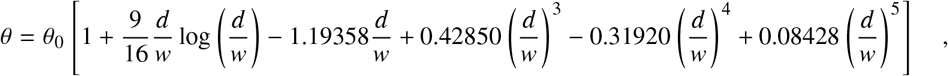

where *θ*_0_ is the diffusion permeability for a molecule of vanishingly small size, which would remain unaffected by the restricted diffusion. Assuming that the observed changes in diffusion permeability during SD are caused by a fairly uniform narrowing of the characteristic average width of gaps separating the individual brain cells, it is then possible to estimate these dynamically changing gaps at any desired time from the measured diffusion permeabilities. Because we measured *θ* before the SD induction for two different molecules, the TMA with *d* = 0.5 nm, and dex3 with *d* = 2.8 nm, we can divide these two diffusion permeabilities to eliminate *θ*_0_. Numerically extracted pore width in propionate before the SD, *w* ≈ 12.6 nm, is subsequently substituted back to calculate *θ*_0_ ≈ 0.475. Under our simplifying assumption of a uniform narrowing of *w*, *θ*_0_ would remain almost constant during the SD event (8). We can thus obtain an estimate of characteristic pore width w at any time. E.g., when the diffusion of dex3 stops at the peak of SD, we get *w* ≈ 2.8, equal to the dex3 hydrodynamic diameter, as would be expected. Interestingly, Thorne and Nicholson (1) estimated that ECS channels modeled as sheet-like pores narrow down to 3.2 nm during terminal ischemia that shares many similarities with spreading depression (24). During the recovery, *θ* = 0.105 and *w* ≈ 5.3, and one hour later, *θ* = 0.239 and *w* ≈ 10.9. Note that the extracellular volume fraction *α* would follow the same course, as long as the changes in ECS volume were primarily due to the approximately uniform changes in pore width *w*. This new indirect approach can thus furnish an estimate for the dynamic behavior of the extracellular volume fraction parameter, not normally accessible by the IOI method. The RTI method can obtain much more reliable estimates of ECS volume fraction but it requires an environment stable for at least tens of seconds. Another alternative method for assessing changes in the ECS volume is to monitor ECS concentration of some background molecule, e.g., TMA, with an ion-selective microelectrode sampling a single spatial location (45). However, this approach is limited to very fast or very homogeneous changes when diffusion of the background molecule in the vicinity of the sampling location can be neglected.

Clinically, SD is thought to underlie a visual aura that precedes the migraine headaches (20–22). Certain SD-like phenomena are also observed in ischemia, trauma, and during seizures (23–25). It is also important to consider long-term consequences of SD and SD-like events on brain well-being. Decreased diffusion permeability for macromolecules and significant reduction of ECS volume during SD and SD-like events will increase concentrations of endogenous macromolecules trafficked in brain ECS. These substances are typically larger than dex3 studied here and therefore more affected; this can have devastating consequences for brain health. At one end of the spectrum are proteins like amyloid-*β*, tau and *α*-synuclein which have physiological functions but cannot be allowed to accumulate (46, 47). At the other end of spectrum are the growth factors which could play a role in cancer and epilepsy (48, 49). In both cases, recurring or long-lasting decreases in diffusion permeability and ECS volume would increase the concentration of these proteins beyond normal physiological range, which may facilitate neurodegeneration and brain disorders. Indeed, epidemiological studies indicate that medical history of migraine correlates with cognitive decline and increased risk of dementia at older age (50, 51).

## CONCLUSION

This study has two outcomes. First, we developed the TR-IOI method to detect extracellular diffusion of fluorescent molecules with high temporal resolution. In current study, dextran (MW 3000) labeled with Texas Red was used and the time resolution was 1–2 s. If a smaller and faster marker molecule was used, the temporal resolution could be improved even further. The TR-IOI can employ a variety of fluorescent molecules and can be applied to the brain in conditions when fast transient changes in ECS occur, such as during SD, oscillations, or epileptic activity. Second, we demonstrated the utility of this method for the SD phenomenon where changes in extracellular diffusion were reported previously. We discovered a transient trapping of a small dextran molecule in the ECS of rat hippocampal slices during the SD.

## AUTHOR CONTRIBUTIONS

JH and SH designed the research. JH carried out theoretical and programming work while SH carried out experimental work. JH and SH wrote the article.

## ACKNOWLEDGMENTS

This work was supported by NIH grants R56-NS047557 and R01-NS047557 (PI S. Hrabetova). Parts of this work have previously been reported in the form of conference abstracts.

